# Nonsense suppression position effect implicates poly(A)-binding protein in the regulation of translation termination

**DOI:** 10.1101/793158

**Authors:** Chan Wu, Bijoyita Roy, Feng He, Allan Jacobson

**Affiliations:** Department of Microbiology and Physiological Systems, University of Massachusetts Medical School, 368 Plantation Street Worcester, MA 01655

**Keywords:** Nonsense codon, PTC, readthrough, poly(A), Pab1

## Abstract

Readthrough of translation termination codons, also known as nonsense suppression, is a relatively inefficient process mediated by ribosomal A site recognition and insertion of near-cognate tRNAs. Multiple factors influence readthrough efficiency, including nonsense codon specificity and context. To determine whether nonsense codon position in a gene influences the extent of readthrough, we generated a series of *LUC* nonsense alleles and quantitated both readthrough and termination efficiencies at each nonsense codon in yeast cells lacking nonsense-mediated mRNA decay (NMD) activity. Readthrough efficiency for premature termination codons (PTCs) manifested a marked dependence on PTC proximity to the mRNA 3’-end, decreasing progressively across the *LUC* ORF but increasing with 3’-UTR lengthening. These effects were eliminated, and translation termination efficiency decreased considerably, in cells harboring *pab1* mutations. Our results support a critical role for poly(A)-binding protein in the regulation of translation termination and suggest that inefficient termination is the trigger for NMD.

## INTRODUCTION

Translation termination in eukaryotes is orchestrated by the release factors eRF1 and eRF3 when any of the three nonsense codons (UAA, UAG, or UGA) in an mRNA occupies the A site of a ribosome (Alkalaeva et al., 2006; Stansfield et al., 1995; Zhouravleva et al., 1995). eRF1 recognizes A site-localized nonsense codons and hydrolyzes peptidyl-tRNA, whereas eRF3 interacts with eRF1 and stimulates termination via its GTPase activity (Alkalaeva et al., 2006; Salas-Marco and Bedwell, 2004). eRF1 and near-cognate (nc) tRNAs, i.e., tRNAs capable of base-pairing with nonsense codons at two of the three standard codon positions, compete for binding to the ribosomal A site (Brown et al., 2015). Although this competition is inefficient, when successful it leads to insertion of an amino acid and continuation of translational elongation until the next in-frame nonsense codon is encountered (Roy et al., 2016; Roy et al., 2015a). Such a bypass of translation termination is designated nonsense suppression or readthrough (Brenner et al., 1965; Keeling et al., 2014; Peltz et al., 2013).

Multiple factors appear to influence the competition between nc-tRNAs and eRF1, and hence the extent of nonsense suppression, including: a) the identity of the stop codon, with UGA promoting higher levels of readthrough than UAG or UAA (Howard et al., 2000; Loughran et al., 2014; Manuvakhova et al., 2000), b) the immediate context of a stop codon, with the highest readthrough levels typically occurring when adenine precedes and cytosine follows the stop codon (Bonetti et al., 1995; McCaughan et al., 1995; Mottagui-Tabar et al., 1998; Tork et al., 2004), c) specific sequences or structures 3’ to the stop codon (Anzalone et al., 2019; Cridge et al., 2018; Harrell et al., 2002; Namy et al., 2001; Skuzeski et al., 1991), and d) the integrity of the release factors and the ribosome (Carnes et al., 2003; Chernoff et al., 1994; Liu and Liebman, 1996; Loenarz et al., 2014; Serio and Lindquist, 1999; Singleton et al., 2014; Velichutina et al., 2000). Another important determinant of readthrough efficiency may be the relative position of the stop codon within the open reading frame (ORF). Toeprinting experiments in yeast demonstrated that termination at premature termination codons (PTCs) is considerably slower than that at normal termination codons (NTCs) (Amrani et al., 2004). This difference predicts that readthrough should occur more readily at PTCs than at NTCs, an expectation borne out by results obtained when cultured cells, animals, or patients are treated with the readthrough-promoting drug ataluren (Hirawat et al., 2007; Welch et al., 2007) as well as by recent studies of non-canonical genetic codes in Ciliates (Heaphy et al., 2016; Swart et al., 2016; Zahonova et al., 2016).

These functional differences between PTCs and NTCs are consistent with the notion that the position of a PTC within an ORF may influence its susceptibility to readthrough and suggest that PTCs closest to the 3’-end of an ORF, i.e., those most likely to have NTC-like contexts, may be the least likely to have high efficiencies of readthrough whereas those more 5’-proximal are likely to have elevated readthrough efficiencies. Here, we have investigated the role of ORF position of PTCs in the regulation of readthrough in yeast. We constructed PTCs at six different positions of the firefly *LUC* ORF, expressed these alleles in yeast cells inactive for the nonsense-mediated mRNA decay (NMD) pathway, and assessed both readthrough and termination efficiencies at each ORF position by quantitating both the full-length protein and the premature termination products expressed from each PTC allele. Our results revealed a position-dependent effect for readthrough and termination efficiencies, and suggested that nonsense codon proximity to the mRNA 3’-end was an important determinant of termination efficiency. Consistent with earlier studies implicating a role for poly(A)-binding protein in the regulation of translation termination (Ivanov et al., 2016) we find that deletion of the yeast *PAB1* gene markedly reduces premature termination efficiency, thus yielding substantial enhancement of PTC readthrough.

## RESULTS

### *LUC* reporters for readthrough assays

To determine the extent of nonsense codon readthrough at different positions of an ORF we constructed a set of reporter genes with PTCs positioned at six locations within the firefly luciferase (*LUC*) coding region (Figure 1A). Each of these constructs has a stop codon at a designated position, a 3X-HA tag at the ORF N-terminus to facilitate the detection of premature termination products, and a StrepII-FLAG (SF) tag at the ORF C-terminus to facilitate the detection of full-length readthrough protein. Each individual *LUC* cassette was cloned into the pRS315 yeast centromere vector, flanked by the promoter region, 5’-UTR, and 3’-UTR of the yeast *TPI1* gene, and then transformed into WT and *upf1Δ* cells to assess expression of the Luc protein. To increase the generation of full-length readthrough products, we used a weak terminator (UGA CAA) (Bonetti et al., 1995) for each PTC reporter while including a strong terminator (UAA) in the normal termination codon (NTC) position. As controls, an NTC reporter (lacking any nonsense codons in the coding region) and an empty vector were also transformed into WT and *upf1Δ* cells.

**Figure 1.**
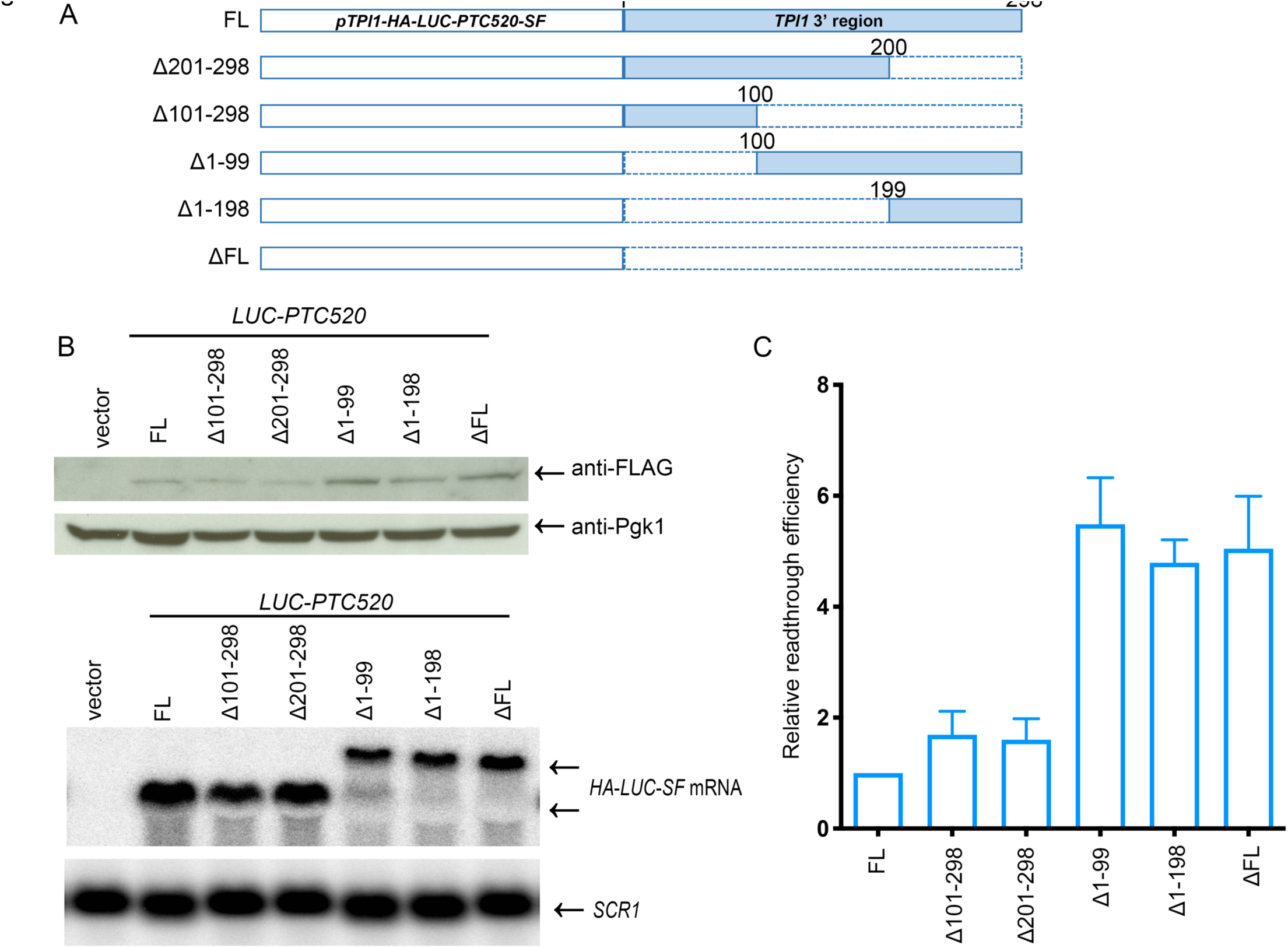
PTC readthrough efficiency decreases across the *LUC* ORF in *upf1Δ* cells A. PTC readthrough reporters. Schematic representation of the respective positions of PTCs and the normal termination codon (NTC) within individual *HA-LUC-SF* constructs used to analyze readthrough and termination efficiency in yeast. The UGACAA sequence context was used in all PTC alleles, each of which was transformed into WT or *upf1Δ* cells to assess readthrough and termination efficiencies. HA, S, F= HA, StrepII, and FLAG epitope tags, respectively. **B. Western analyses of full-length readthrough proteins and prematurely terminated polypeptides expressed from *LUC* PTC alleles.** Each of the six *LUC* PTC alleles, a vector control, and a wild-type *LUC* allele was expressed in separate cultures of WT or *upf1Δ* yeast cells that were subsequently harvested, lysed, and analyzed by western blotting using anti-FLAG antibodies to detect full-length readthrough products, anti-HA antibodies to detect premature termination products, and anti-Pgk1 antibodies to detect the Pgk1 control protein. The arrow in the upper panel indicates the full-length readthrough protein and the asterisks in the middle panel indicate premature termination products. Figure S1D shows the complete blot used for the top panel of Figure 1B. **C. Northern analyses of *LUC* mRNA levels in cells expressing different PTC alleles.** Northern blotting and phosphorimaging were utilized to measure the level of *LUC* mRNA and *SCR1* RNA in each sample. **D, E. Relative readthrough efficiency at each of the six *LUC* PTCs.** Densitometry was utilized to quantitate the western blots in part B and the northern blots in part C. Relative readthrough efficiency of each sample was calculated as the ratio of full-length Luc protein level (anti-FLAG) normalized to Pgk1 level divided by *LUC* mRNA level normalized to the level of *SCR1* RNA. The efficiency of *LUC* PTC20 expression in WT cells (in part D) or *upf1Δ* cells (in part E) was set as 1. Three independent experiments were performed and the results shown are the average of all three +/− SEM. 1% Luc-wt and 2% Luc-wt: aliquots of extracts from cells expressing a wild-type *LUC* allele that were 1/100^th^ and 1/50^th^ the volume of samples from cells expressing PTC alleles.

### PTC readthrough efficiency decreases across the *LUC* ORF

To compare the efficacy of PTC readthrough at different positions of the *LUC* ORF, cells harboring the respective *LUC* alleles were grown, harvested, and analyzed in parallel. Relative readthrough efficiencies (Figures 1D, E) were determined by normalizing western blotting results with anti-FLAG antibodies (Figure 1B, upper panel) to the level of an internal control protein (Pgk1) in the same samples (Figure 1B, lower panel) and to the level of *LUC* mRNA in each sample (determined by northern blotting, and also normalized to a control RNA, *SCR1*; Figure 1C). In WT cells, the levels of full-length readthrough protein obtained from the early PTC alleles, PTC20, PTC41, and PTC160, were comparably low, whereas those obtained from the middle PTCs, PTC386 and PTC460, were approximately 1.5-to 2-fold higher (Figure 1B, left, upper panel; Figure 1D, left panel). That obtained with the late PTC, PTC520, was the lowest of the set. Every PTC allele in WT cells showed readthrough levels of luciferase that were less than 1% of the level obtained from the WT *LUC* allele (Luc-wt) (Figure 1B, upper panel).

Upf1 is a key regulator of NMD, and deletion of its gene not only stabilizes nonsense-containing mRNAs, but also enhances nonsense codon readthrough and inactivates the degradation of prematurely terminated polypeptides (He and Jacobson, 2015; Johansson and Jacobson, 2010; Kuroha et al., 2009; Leeds et al., 1991). Accordingly, to augment detection of readthrough products, and to minimize mRNA variability as a potential complication, we repeated the same experiments in *upf1Δ* cells. The extent of readthrough in *upf1Δ* cells was highest when a PTC was proximal to the N-terminus of the *LUC* ORF (PTC20 and PTC41 showed ~2% of Luc-wt) and diminished progressively as the PTC positions approached the C-terminus of the ORF (Figure 1B, upper panel). After normalizing to *LUC* mRNA levels (Figure 1C), the relative readthrough efficiencies of PTCs in *upf1Δ* cells showed a clear position effect in which readthrough efficiency decreased across the *LUC* ORF (Figures 1D, E). These variations in readthrough efficiency were not likely to be attributable to differences in the stability of the respective full-length readthrough proteins (Kuroha et al., 2009) because treating the cells with the proteasome inhibitor MG132 did not alter the trend of the readthrough proteins across the ORF (Figures S1A, B).

In parallel with our western blotting assays for full-length readthrough proteins we also employed western blotting with anti-HA antibodies to detect prematurely terminated polypeptides originating from each PTC allele. These products were readily detected from PTC160, PTC386, PTC460, and PTC520 (Figure 1B, middle panel, and Figure S1A, middle panel; see red asterisks). Corrected for the Pgk1 control protein and mRNA levels, the recovery of prematurely terminated polypeptides in *upf1Δ* cells showed a general upward trend from PTC160 to PTC460, a result consistent with our observation of decreased readthrough across the same region of the *LUC* ORF (Figure S2A and B, left panels). Further, our inability to detect any premature termination products from the PTC20 and PTC41 alleles expressed in WT and *upf1Δ* cells was not likely to be due to their failure to be recovered by western blotting. This conclusion follows from an experiment in which we synthesized a polypeptide containing the 3X-HA epitope and the first 19 amino acids of luciferase (i.e., the polypeptide that would be generated by premature termination at PTC20), mixed it with a lysate from cells expressing PTC20, and subjected the sample to our western blotting procedure. Figure S1C shows that this peptide could be detected.

### Alterations of *LUC* mRNA 3’-UTR lengths correlate with changes in relative readthrough efficiencies

Earlier studies have suggested that interactions between mRNA-associated poly(A)-binding protein (Pab1 in yeast) and the release factor eRF3 enhance translation termination efficiency (Amrani et al., 2004; Heaphy et al., 2016; Ivanov et al., 2016; Roque et al., 2015; Swart et al., 2016; Zahonova et al., 2016). To test whether the progressive reduction in *LUC* PTC readthrough efficiencies we observed in *upf1Δ* cells reflects changes in PTC proximity to Pab1 (or any other 3’-UTR-localized regulatory factor), we began with the reporter manifesting the least efficient readthrough (*LUC-PTC520*) and generated a set of deletions in the *TPI1* fragment of that gene that encodes its 3’-UTR (Figure 2A). The reporters harboring these deletions were then transformed into *upf1Δ* cells and the effects of the deletions on PTC readthrough efficiencies were determined as in Figure 1. Western blotting with anti-FLAG antibodies indicated that the level of full-length readthrough protein obtained from the Δ101-298 and Δ201-298 alleles remained similar to that of the *LUC-PTC520* allele, but increased in the samples obtained from the Δ1-99, Δ1-198, and ΔFL alleles (Figure 2B, upper panel). After normalization to the Pgk1 control protein and the mRNA levels of each sample, we saw a 5-6-fold increase in the relative readthrough efficiencies of the mRNAs derived from the Δ1-99, Δ1-198, and ΔFL alleles (Figure 2C).

**Figure 2.**
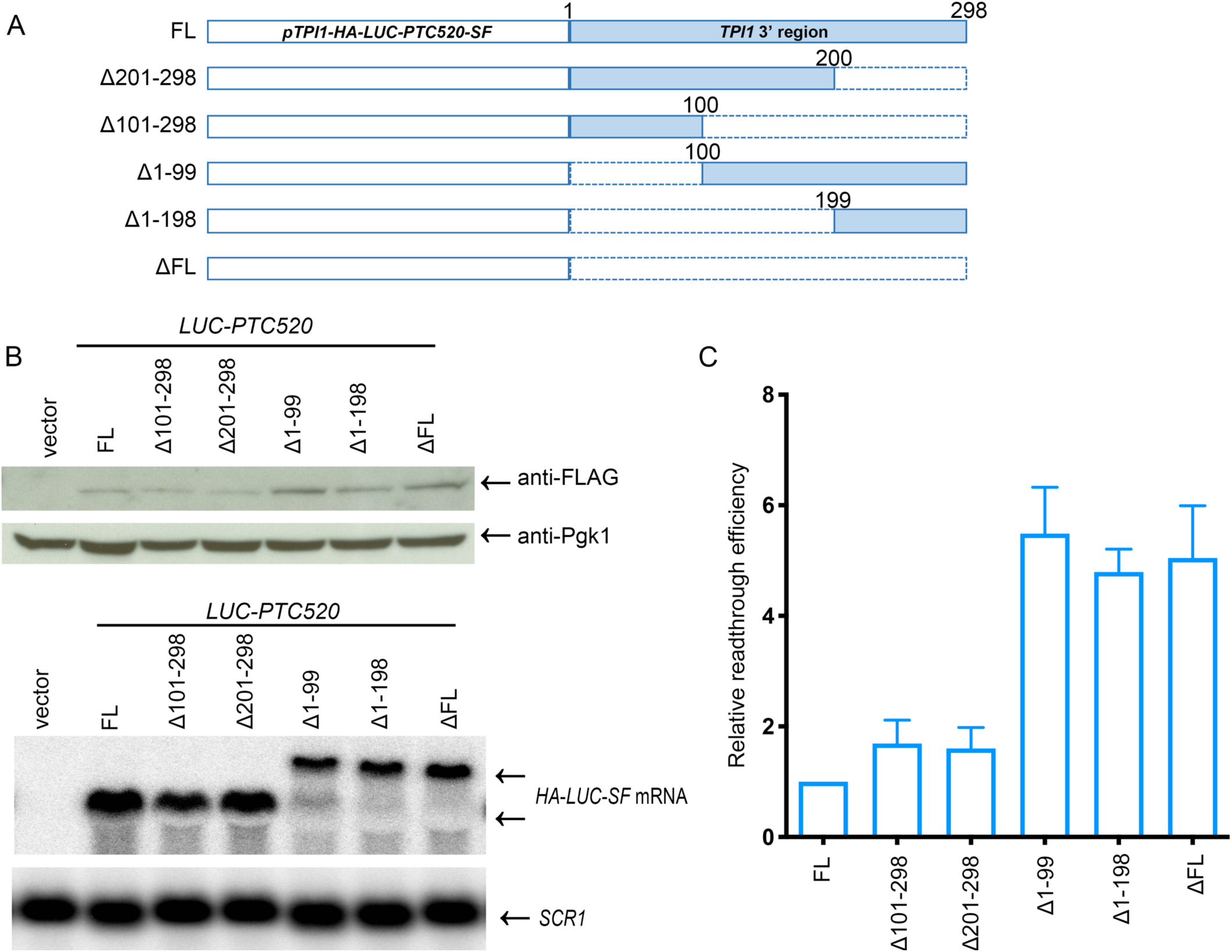
Alterations of the 3’-UTR associated with the *LUC* ORF affect readthrough efficiency. **A. Schematic representation of 3’-UTR deletion reporters.** *LUC PTC520* alleles harboring different deletions (indicated by the dashed bars) in the *TPI1* 3’ region were constructed and each of the these alleles was separately transformed into *upf1Δ* cells to assess expression of both mRNA and protein. *pTPI1*=*TPI1* promoter. *SF*=*StrepII*-*FLAG* tag. FL=Full-length *TPI1* 3’-region. **B. Western and northern analyses of 3’-UTR deletion reporters.** Yeast *upf1Δ* cells harboring the original *LUC* PTC520 allele, each of the 3’-UTR deletion alleles, and an empty vector were cultured separately and samples were processed as in Figure 1. In the upper panel (western blot), anti-FLAG antibodies detected full-length readthrough products and anti-Pgk1 antibodies detected the control protein in each sample. In the lower panel, northern blotting was used to assess the levels of *LUC* mRNA and *SCR1* RNA in each sample. **C. Relative readthrough efficiency of 3’-UTR deletion reporters.** Relative readthrough efficiency was calculated as in Figure 1D, using the data obtained from western and northern blotting experiments like those in Figure 2B. The results shown are the average of three independent experiments +/− SEM.

Northern blotting of the mRNAs expressed by each of the 3’-UTR deletion alleles revealed that the transcripts generated by the Δ1-99, Δ1-198, and ΔFL alleles were significantly longer than that derived from the original *LUC-PTC520* allele (Figure 2B lower panel), suggesting that these alleles are commonly defective in *TPI1* 3’-UTR-mediated cleavage/polyadenylation and most likely use alternative cryptic polyadenylation signals in the downstream vector sequences for mRNA formation. Consistent with this idea, our 3’-RACE and sequencing analyses indicated that transcripts generated from the *LUC-PTC520* allele terminate within the *TPI1* 3’ fragment at a single site that is 75 nt downstream of the *LUC* ORF, whereas transcripts generated from the Δ1-99, Δ1-198, and ΔFL alleles all terminate within the vector at a site that is more than 600 nt downstream of the *LUC* ORF (data not shown). To determine whether the observed enhancement of readthrough by the Δ1-99, Δ1-198, and ΔFL alleles is due to the increased length of their 3’-UTRs or to loss of putative positive 3’-UTR regulatory elements involved in translation termination we constructed an additional set of *LUC-PTC520* reporters that harbored small deletions in the gene’s 3’-UTR (Figure 3A). These alleles were transformed into *upf1Δ* cells and their expression was monitored by western and northern blotting as above. The two *LUC*-*PTC520* alleles harboring small deletions in the *TPI1* 3’-UTR, Δ1-24 and Δ25-50, produced mRNAs similar in length to that from the original *LUC*-*PTC520* allele, and these two alleles also had similar levels of full-length readthrough products and readthrough efficiencies as the *LUC*-*PTC520* allele (Figures 3B and C). In contrast, the *LUC-PTC520* allele harboring a large deletion in the *TPI1* 3’-UTR, Δ1-50, produced mRNAs that included a longer isoform than that from the *LUC*-*PTC520* allele, and this allele had a significantly higher level of full-length readthrough product and readthrough efficiency than the *LUC*-*PTC520* allele (Figures 3B and C). Since the increased readthrough efficiency of the Δ1-50 allele was not matched by either the Δ1-24 or the Δ25-50 allele (Figure 3C) it is likely that the principal readthrough-enhancing consequences of the Δ1-50 deletion are attributable to 3’-UTR lengthening.

**Figure 3.**
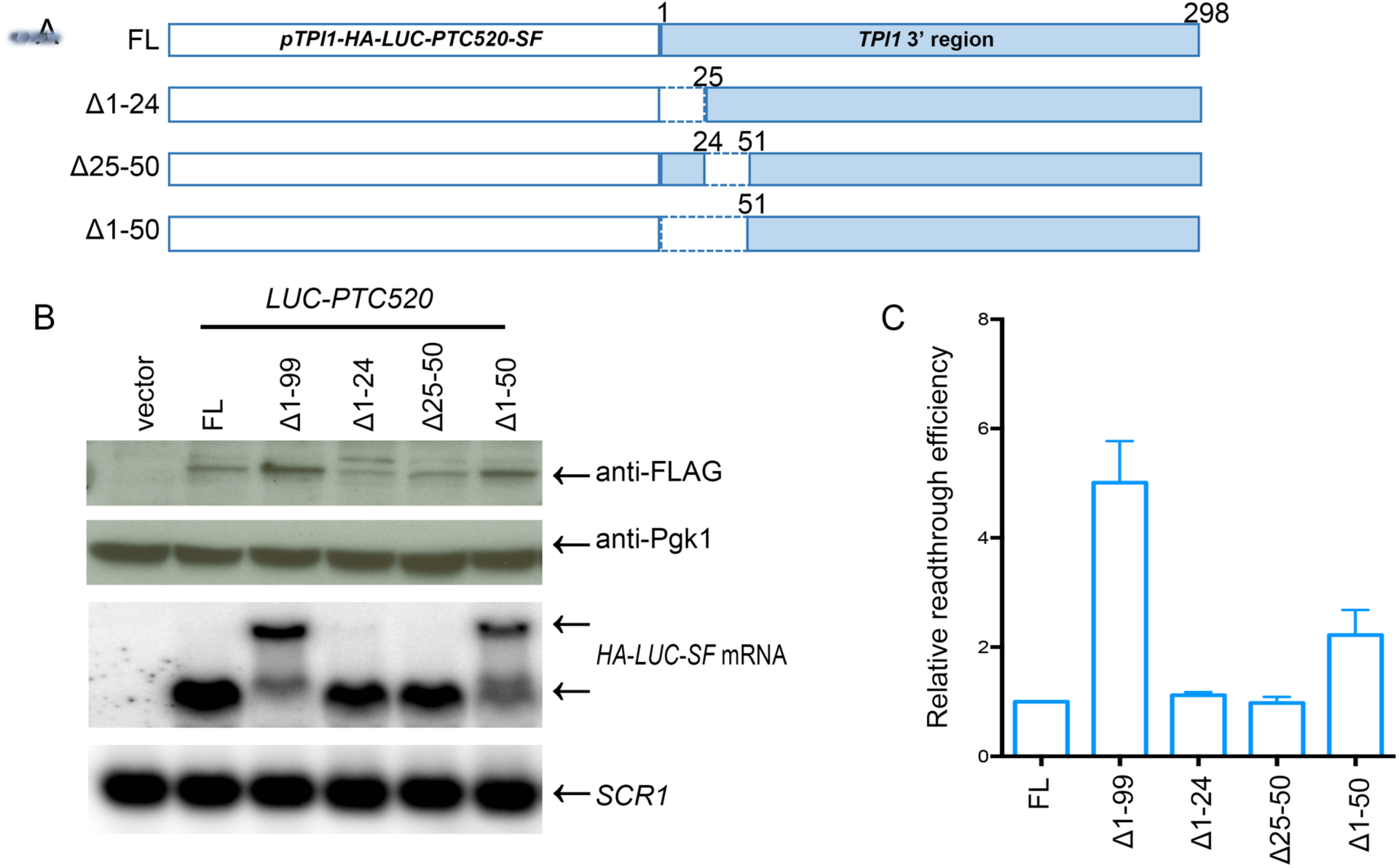
3’-UTR deletions only affect *LUC* PTC readthrough efficiency when mRNA length is altered. **A. Schematic representation of *LUC* PTC520 reporters with different deletions in the *TPI1* 3’-UTR.** A set of *LUC* PTC520 alleles harboring different deletions in the *TPI1* 3’-UTR (indicated by dashed bars) was constructed. As in Figure 2, each of the constructs was transformed into *upf1Δ* cells for subsequent determination of *LUC* mRNA and protein levels. *pTPI1*=*TPI1* promoter. *SF*=*StrepII*-*FLAG* tag. FL=Full-length *TPI1* 3’-region. **B. Analyses of mRNA and readthrough products generated from *LUC* PTC520 reporters with different 3’-UTR deletions.** In the upper panel, anti-FLAG antibodies were used to detect full-length readthrough protein generated from each of the alleles, and anti-Pgk1 antibodies were used to detect the control Pgk1 protein. In the lower panel, northern blotting was used to assess the levels of *LUC* mRNA and *SCR1* RNA in each sample. **C. Relative readthrough efficiency of *LUC* PTC520 reporters with different 3’-UTR deletions.** Relative readthrough efficiencies were calculated as in Figure 1D. The results shown are the average of three independent experiments +/− SEM.

### 3’-UTR length, but not a specific sequence element, regulates the efficiency of PTC readthrough

As noted above, increased PTC readthrough efficiencies of *LUC*-*PTC520* reporters appear to correlate with lengthened 3’-UTRs. To further address this possibility, we carried out two additional experiments. In the first, we tested whether shortening of a long 3’-UTR by inserting a strong polyadenylation signal upstream of the cryptic polyadenylation site can diminish the PTC readthrough efficiency associated with that transcript. Based on the Δ1-99 allele from Figure 2, we constructed two *LUC-PTC520* alleles that lack the original *TPI1* 3’-UTR, but contain different sequences from the *TPI1* 3’ region (i.e., from nt 100 to 199 or 100 to 298) with a strong polyadenylation signal from the *GAL7* gene (Bucheli, He et al. 2007) inserted within the *TPI1* 3’ sequences (Figure 4A). We designated these two constructs as S and L alleles and expected each of these alleles to produce transcripts with 3’-UTR lengths of about 75 nt. The S and L 3’-UTR reporters were transformed into *upf1Δ* cells and western and northern analyses were used to monitor their expression as above. Northern blotting showed that the size of the *LUC* mRNAs generated from both alleles was now comparable to that derived from the original *LUC*-*PTC520* allele (Figure 4B). Further, the elevated PTC readthrough efficiency resulting from a lengthened 3’-UTR (Δ1-99 allele) was reduced to the level of the *LUC-PTC520 FL* allele (Figure 4B, C).

**Figure 4.**
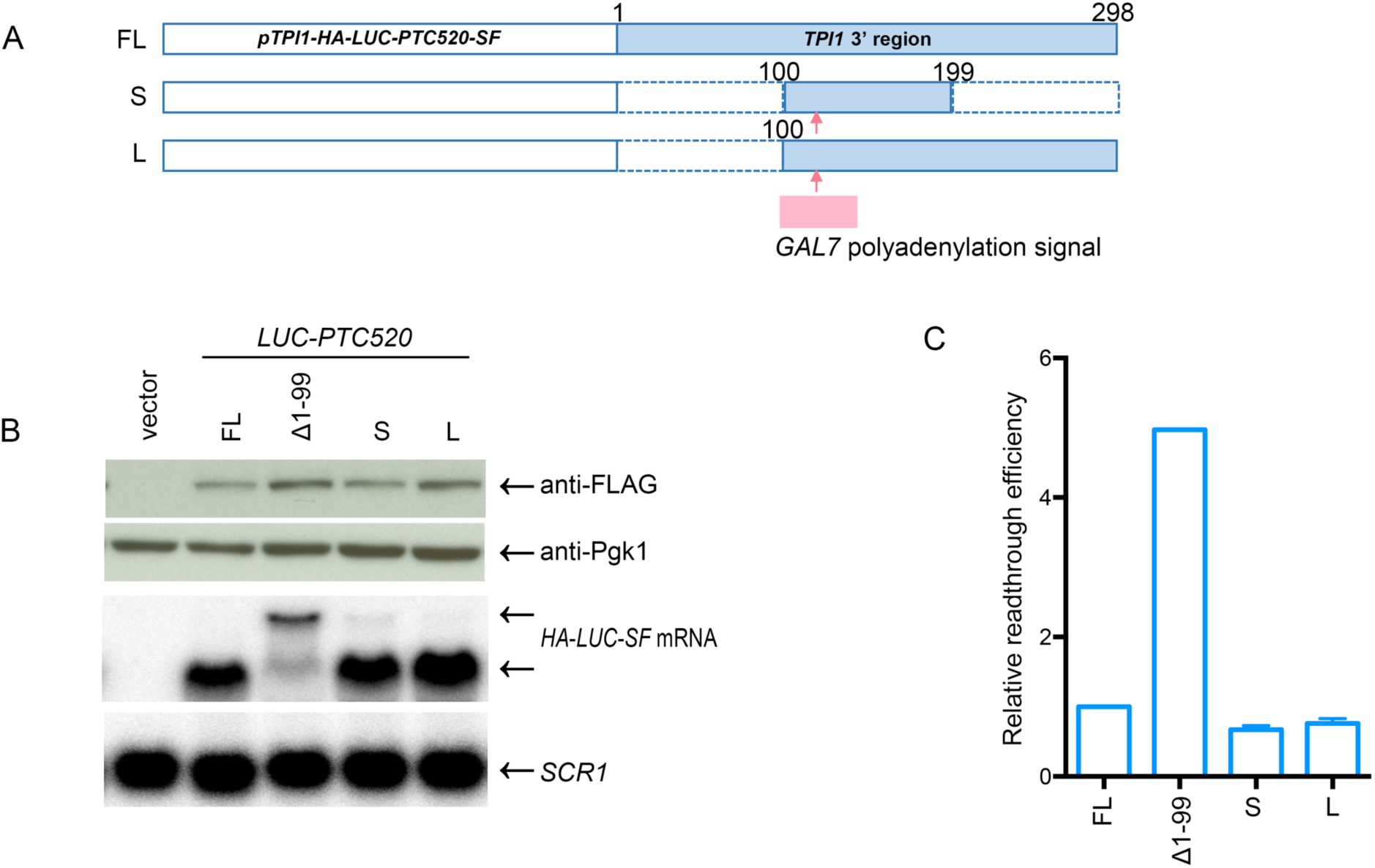
The increased readthrough efficiency of mRNAs with extended 3’-UTRs is reversed when the 3’-UTR is shortened. **A. *LUC* PTC520 alleles with insertions of the *GAL7* cleavage/polyadenylation signal.** The *GAL7* cleavage/polyadenylation signal (shown in pink) was inserted between nts 130 and 131 of short (100nt-199nt) and long (100nt-298nt) derivatives of the *TPI1* 3’ region to generate the S and L alleles. These *LUC* PTC520 alleles were transformed into *upf1Δ* cells to assess mRNA and protein expression. *pTPI1*=*TPI1* promoter. *SF*=*StrepII*-*FLAG* tag. FL=Full-length *TPI1* 3’-region. S, L= short or long derivatives of the *TPI1* 3’-UTR. **B. mRNAs and readthrough products generated from the *LUC* PTC520 S and L alleles.** As in Figure 1B, the levels of full-length readthrough protein and Pgk1 control protein derived from cells expressing each allele were analyzed by western blotting. In the lower panel, northern blotting was used to assess each sample’s levels of *LUC* mRNA and *SCR1* RNA. **C. Relative readthrough efficiencies of the *LUC* PTC520 S and L alleles.** Relative readthrough efficiencies were calculated as in Figure 1D. The results shown are the average of three independent experiments +/− SEM. In both B and C, results from expression of the *LUC* PTC520 FL and Δ1-99 alleles were included for comparison.

In the second experiment we tested directly the effects of 3’-UTR length on PTC520 readthrough efficiency. We made use of the *LUC PTC520 ΔFL* allele from Figure 2 and constructed a series of *LUC PTC520* alleles that lacked a *TPI1* 3’-UTR by inserting the *GAL7* polyadenylation signal at different positions in the vector sequences downstream of the *LUC* ORF (Figure 5A). Transcripts generated from these *LUC PTC520* alleles lack any *TPI1* 3’-UTR sequences, but contained different sequences in their 3’-UTRs with lengths ranging from 60 to 600nt. Quantitation of the expression of these alleles in *upf1Δ* cells showed that the level of full-length readthrough protein and the relative readthrough efficiencies increased progressively with 3’-UTR length (Figures 5B and C). Collectively, the results of Figures 2-5 indicate that the length of the 3’-UTR associated with the *LUC* mRNA plays an important *cis*-acting role in regulating translational readthrough of PTCs in yeast.

**Figure 5.**
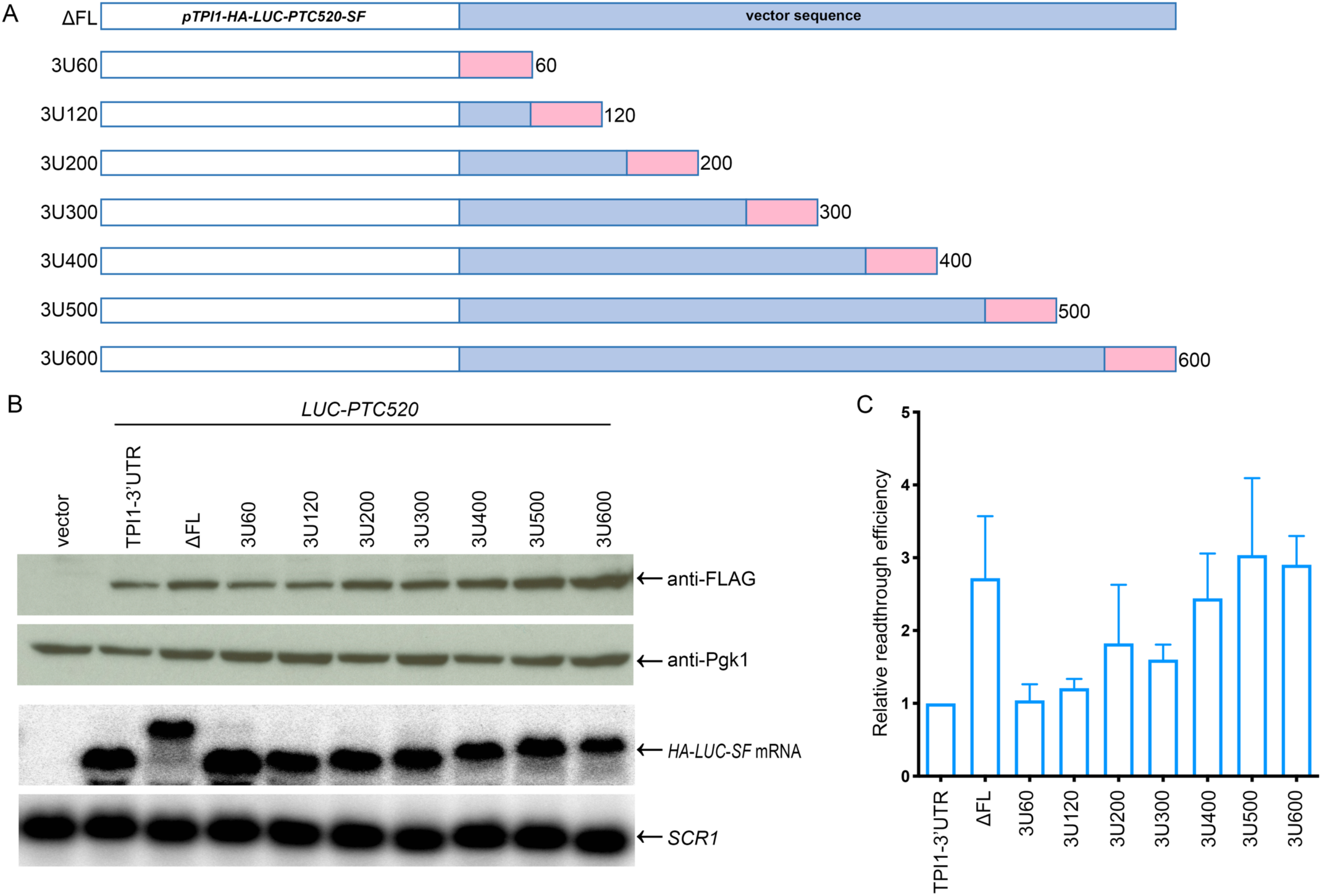
The efficiency of readthrough of *LUC* PTC520 mRNA varies directly with the length of its associated 3’-UTR. **A. Schematic representation of *LUC* PTC520 alleles with defined 3’-UTR lengths.** The *GAL7* cleavage/polyadenylation signal (pink rectangles) was inserted at different locations downstream of the *LUC* PTC520 ORF (ΔFL allele in Figure 2) to generate a set of *LUC* PTC520 alleles. The individual alleles were transformed into *upf1Δ* cells for assessment of *LUC* mRNA and protein expression. mRNAs generated from this set of alleles have 3’-UTR lengths ranging from 60 to 600 nt. *pTPI1*=*TPI1* promoter. *SF*=*StrepII*-*FLAG* tag. FL=Full-length *TPI1* 3’-region. **B. Analyses of the mRNAs and readthrough products generated by *LUC* PTC520 alleles with defined 3’-UTR lengths.** *LUC* PTC520 alleles harboring 3’-UTRs of 60bp, 120bp, 200bp, 300bp, 400bp, 500bp, or 600bp were expressed in *upf1Δ* cells and lysates of the respective cells were analyzed by western blotting and northern blotting as in Figure 1B. **C. Relative readthrough efficiencies of *LUC* PTC520 alleles with 3’-UTRs of defined lengths.** Relative readthrough efficiencies were calculated as in Figure 1D. The results shown are the average of four independent experiments +/− SEM.

### Poly(A) binding protein restricts PTC readthrough and maintains the PTC position effect

In light of the correlation between 3’-UTR length and PTC readthrough seen in Figures 2–5 we sought to test directly whether the observed readthrough phenotypes reflected PTC proximity to mRNA-associated Pab1 protein. *PAB1* is an essential gene in yeast (Sachs et al., 1986; Sachs et al., 1987), but its absence can be studied in several suppressor mutants. Pbp1 was identified as a Pab1-interacting protein and *pbp1Δ* cells suppress a *pab1Δ* allele without having significant effects on mRNA translation or decay (Kato et al., 2019; Mangus et al., 1998a; Yang et al., 2019). Accordingly, we generated *pab1Δ pbp1Δ upf1Δ* yeast cells and transformed them with the *pRS316*-*LUC*-*PTC* reporters and control plasmids equivalent to those analyzed in the experiments of Figure 1. As additional controls, each of these reporters was also transformed into *pbp1Δ upf1Δ* cells. Readthrough efficiencies of the *LUC*-*PTC* reporters were determined in *upf1Δ*, *pbp1Δ upf1Δ*, and *pab1Δ pbp1Δ upf1Δ* cells and the results of these analyses are shown in Figure 6. Consistent with previous conclusions about the modest impact of the *pbp1Δ* allele on mRNA translation and decay, we found the relative readthrough efficiencies of *LUC*-*PTC* reporters in *pbp1Δ upf1Δ* cells to be quite similar to those seen in *upf1Δ* cells (Figure 6C). However, in *pab1Δ pbp1Δ upf1Δ* cells the relative readthrough efficiencies of all *LUC*-*PTC* reporters were much higher than in *upf1Δ* or *pbp1Δ upf1Δ* cells (Figure 6C), with readthrough efficiencies of PTC20, PTC41, PTC160, PTC386, PTC460, and PTC520 in *pab1Δ pbp1Δ upf1Δ* cells approximately 2-to 8-fold higher than those observed in the two other strains. Whereas readthrough efficiencies across the *LUC* ORF in *upf1Δ* cells continuously decreased as the ribosome progressed from 5’ to 3’, in *pab1Δ pbp1Δ upf1Δ* cells this gradient was absent and was replaced by a bell-shaped distribution of readthrough activity across the ORF. Further, in contrast to the increased levels of the *LUC* readthrough proteins (Figure 6A, upper panel), the premature termination products were much lower in *pab1Δ pbp1Δ upf1Δ* cells than in *upf1Δ* or *pbp1Δ upf1Δ* cells (Figure 6A, middle panel), with several premature termination products beyond detection in *pab1Δ pbp1Δ upf1Δ* cells. Although termination efficiencies of *LUC-PTCs* in *pab1Δ pbp1Δ upf1Δ* cells could not be quantitated, those in *pbp1Δ upf1Δ* cells could and they were found to be 1.5 to 2-fold lower than those in *upf1Δ* cells (Figure S2A).

**Figure 6.**
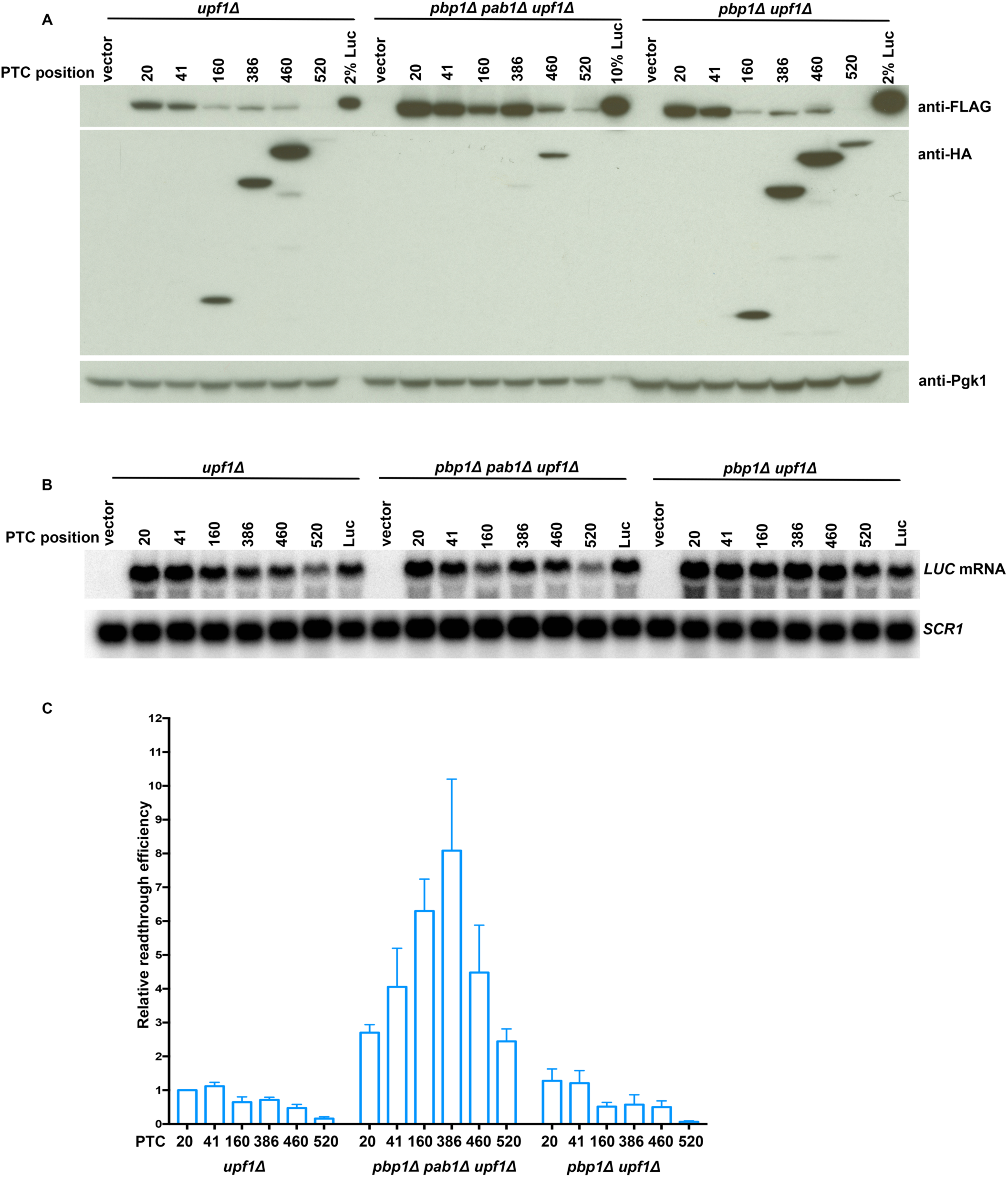
Deletion of *PAB1* enhances translational readthrough and disrupts PTC position effects. **A, B. Western and northern analyses of *LUC-PTC* reporter expression in *upf1Δ*, *pab1Δ pbp1Δ upf1Δ*, and *pbp1Δ upf1Δ* cells.** Each of the six *LUC* PTC alleles, an empty vector control, and a wild-type *LUC* allele was expressed in separate cultures of *upf1Δ*, *pab1Δ pbp1Δ upf1Δ*, and *pbp1Δ upf1Δ* cells that were subsequently harvested, lysed, and analyzed by western and northern blotting as in Figure 1B. **C. Relative readthrough efficiencies of different *LUC* PTC reporters in cells with and without Pab1.** The relative readthrough efficiencies of each of the *LUC* PTC alleles in *upf1Δ*, *pab1Δ pbp1Δ upf1Δ*, and *pbp1Δ upf1Δ* cells were calculated as in Figure 1D. The results shown are the average of three independent experiments +/− SEM. 10% Luc: aliquot of extract from cells expressing a wild-type *LUC* allele that was 1/10^th^ the total proteins of samples from cells expressing PTC alleles.

The results of Figure 6 demonstrate that Pab1 restricts PTC readthrough *in vivo*. To begin to assign that function to specific Pab1 domains we examined the consequences of deleting the C-terminal domain of the protein, a deletion that does not impair yeast cell viability (Sachs et al., 1987). We constructed *pab1ΔC upf1Δ* cells, transformed them with the of *LUC*-*PTC* and control reporters, and evaluated PTC readthrough efficiency in these cells. Figure 7A (top panel) shows that the yields of full-length proteins from all six *LUC*-*PTC* reporters in *pab1ΔC upf1Δ* cells were higher than in *upf1Δ* cells. After correction for protein and mRNA controls (Figures 7A, lower panel, and 7B), the relative readthrough efficiencies of PTCs in *pab1ΔC upf1Δ* cells were 2-to 4-fold higher than those of the same alleles in *upf1Δ* cells (Figure 7C). Readthrough efficiencies in *pab1ΔC upf1Δ* cells were “flat” across the first two-thirds of the *LUC* ORF, but decreased for the last two alleles. Although premature termination products were detectable in *pab1ΔC upf1Δ* cells, their termination efficiencies were lower than those of *upf1Δ* cells or *pbp1Δ upf1Δ* cells (Figure S2B).

**Figure 7.**
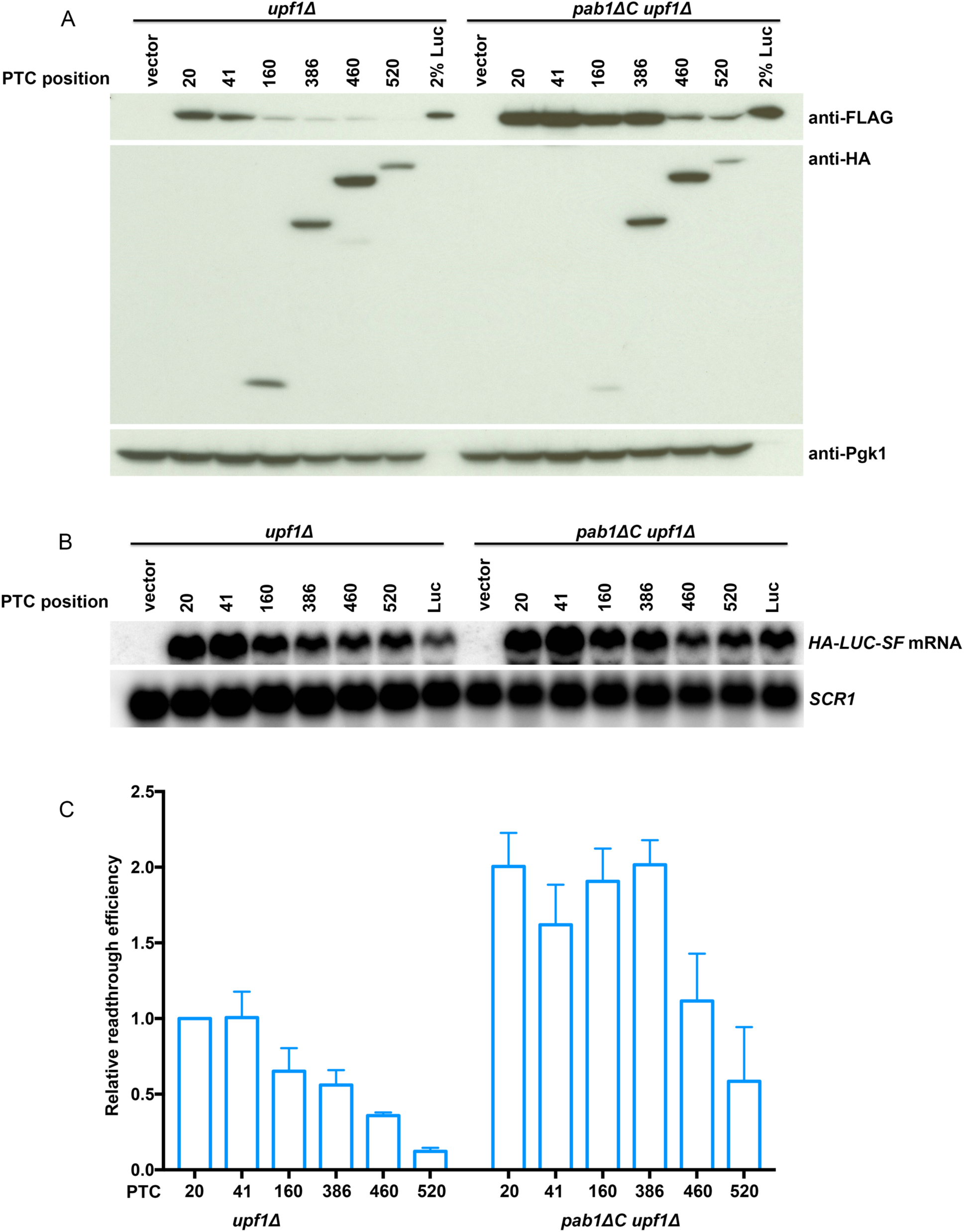
The Pab1 C-terminal domain is required for maintenance of PTC position effects. **A, B. Western and northern analyses of *LUC* PTC reporters in *upf1Δ* and *pab1ΔC upf1Δ* yeast cells.** Each of the six *LUC* PTC alleles, a vector control, and a wild-type *LUC* allele was expressed in separate cultures of *upf1Δ* or *pab1ΔC upf1Δ* cells that were subsequently harvested, lysed, and analyzed by western and northern blotting as in Figure 1B. **C. Relative readthrough efficiencies of *LUC* PTC reporters in *upf1Δ* and *pab1ΔC upf1Δ* cells.** Relative readthrough efficiencies were calculated as in Figure 1D and the results shown are the average of three independent experiments +/− SEM.

## DISCUSSION

### A PTC position effect in the *LUC* ORF is associated with variations in the efficiency of translation termination

In experiments that established the colinearity of genes and their encoded polypeptide chains Sarabhai et al. (1964) also substantiated the prediction of Crick et al. (1961) that nonsense codons promoted the termination of protein synthesis. Initially, the only exceptions to the chain terminating effects of nonsense codons were observed in strains or extracts of *E. coli* expressing suppressor tRNAs (Andoh and Ozeki, 1968; Capecchi and Gussin, 1965; Engelhardt et al., 1965), but it eventually became clear that nonsense codon “readthrough” without suppressor tRNA intervention could be detected in numerous mRNAs of diverse species (Dunn et al., 2013; Eswarappa et al., 2014; Jungreis et al., 2011; Klagges et al., 1996; Loughran et al., 2014; Namy et al., 2003; Namy et al., 2002; Robinson and Cooley, 1997; Schueren et al., 2014; Steneberg and Samakovlis, 2001; Stiebler et al., 2014; Weiner and Weber, 1971). However, even with this considerable history, the mechanisms regulating nonsense codon readthrough remain poorly understood. Here, we demonstrate that in yeast *upf1Δ* cells the position of a PTC in the *LUC* ORF dictates its readthrough efficiency such that a PTC proximal to the 5’-end of the ORF has ~twofold higher readthrough efficiency than that of a PTC proximal to the 3’-end of the ORF (Figures 1B, upper panel, and 1E). Further, in parallel with a consistent reduction in readthrough efficiency from the beginning to the end of the *LUC* ORF, termination efficiency largely appeared to increase (Figures 1B, 6A, 7A, and S2). This position dependence of nonsense codon function is reminiscent of the non-canonical genetic code utilization in some species of protozoa, e.g., *C. magnum*, *Parduczia sp.*, and *Blastocrithidia sp.*, in which all three nonsense codons function as sense codons unless they are located near ORF 3’-ends, where they can still serve their usual termination function (Heaphy et al., 2016; Swart et al., 2016; Zahonova et al., 2016). The decoding of UAA, UAG, and UGA in these atypical genetic codes appears to be mediated in part by mutations that lessen the specificity of the release factor eRF1 and make it an ineffective competitor with tRNAs capable of nonsense codon recognition (Swart et al., 2016). The enhanced specificity of the same eRF1 molecules when ribosomes are translating near the end of ORFs is postulated to arise from proximity to 3’-end-associated poly(A)-binding protein and its stimulatory effects on eRF3, the release factor that normally augments eRF1 function (Swart et al., 2016). As discussed below, poly(A)-binding protein also appears to be a critical regulator of the *LUC* PTC position effects in yeast.

### Yeast Pab1 is a key determinant of the PTC position effect

The results of two different experimental approaches described here both implicate a role for poly(A)-binding protein (Pab1 in yeast) in the regulation of *LUC* PTC readthrough. First, we observed an inverse correlation between the extent of readthrough at a given PTC and the distance of that PTC from the mRNA 3’-end, i.e., the position of the poly(A) tail. This was true for PTCs at different positions of the *LUC* ORF (Figure 1), as well as for PTC520 when it was associated with 3’-UTRs of different lengths (Figures 2-5). Consistent with these readthrough results was a trend toward higher efficiencies of translation termination as PTCs approached the ORF 3’-end (Figures 1B, 7A, S1A, and S2). Collectively, these position effects indicate that the efficiency of termination of *LUC* mRNA translation increases, and readthrough consequently decreases, as a ribosome approaches the mRNA 3’-end. While these results only provide indirect evidence implying a role for Pab1 in termination regulation a second set of experiments provided direct evidence for that role. Figure 6 shows that deletion of the *PAB1* gene leads to large increases in readthrough efficiency at all PTC positions, comparably large reductions in termination efficiency, and loss of the progressive PTC position effects across the *LUC* ORF. Further, the experiments of Figure 6 demonstrate that the *pbp1Δ* mutation required for the maintenance of viability in *pab1Δ* cells does not contribute to these effects.

A role for Pab1 in restricting readthrough is consistent with several earlier studies in multiple systems, including those reporting: a) position dependence of the non-canonical genetic codes in Ciliates (alluded to above) (Heaphy et al., 2016; Swart et al., 2016; Zahonova et al., 2016), b) interactions between eRF3 and PABP from mammalian, *Xenopus*, and yeast cells, and identification of specific interacting domains in the respective proteins, using two-hybrid, co-immunoprecipitation, isothermal titration calorimetry, and surface plasmon resonance methodologies (Cosson et al., 2002a; Cosson et al., 2002b; Hoshino et al., 1999; Hosoda et al., 2003; Jerbi et al., 2016; Kononenko et al., 2010; Roque et al., 2015; Uchida et al., 2002), and c) direct stimulation of peptidyl-tRNA hydrolysis dependent on eRF3:PABP interaction in a reconstituted cell-free system (Ivanov et al., 2016). However, our results differ from those of Roque et al. who found that readthrough efficiency in a dual luciferase assay in yeast decreased when Pab1:eRF3 interaction was interrupted (Roque et al., 2015). In addition to the use of different readthrough reporters a significant difference between our study and that of Roque et al. that may account for this difference was their inclusion of suppressor-tRNA in the strains being tested, i.e., their assays focused on the strong but artificial readthrough signals generated by a cognate suppressor tRNA whereas ours addressed the competition between eRF1 and near-cognate tRNAs that is typical of conventional PTC readthrough (Roy et al., 2016; Roy et al., 2015a).

### The inefficiency of premature termination may define susceptibility to NMD

Although the focus of this study has been the possible role of Pab1:eRF3 interaction in the regulation of translation termination efficiency our results also have important implications for NMD. Earlier work on NMD in yeast showed that: a) PTCs within the first half to two-thirds of an ORF triggered NMD whereas those in the latter part of an ORF had little to no mRNA destabilizing effects (Hagan et al., 1995; Hennigan and Jacobson, 1996, 1997; Peltz et al., 1993; Peltz et al., 1994; Peltz and Jacobson, 1993), b) these PTC position effects in NMD could be mimicked and manipulated by shortening or lengthening mRNA 3’-UTRs (Amrani et al., 2004; Kebaara and Atkin, 2009; Muhlrad and Parker, 1999), and c) NMD could be antagonized by tethering Pab1 or eRF3 proximal to a PTC (Amrani et al., 2004). Indeed, these results, and toeprinting experiments showing that termination at PTCs was considerably slower than that at NTCs (Amrani et al., 2004), led to formulation of the *faux*-UTR model for NMD which postulated that the inefficient termination occurring at PTCs distant from an mRNA’s bound Pab1 allowed for mRNA binding of the Upf factors that trigger NMD (Amrani et al., 2006a; Amrani et al., 2004; Amrani and Jacobson, 2006; Amrani et al., 2006b). Although almost all of the experiments here were carried out in NMD-deficient cells, the position effect of PTCs on NMD susceptibility of the *LUC* mRNA was evident in Figure 1C (compare WT to *upf1Δ*), is consistent with earlier published results, and, together with the data in Figures 1-7, indicates that termination deficiencies at PTCs are independent of Upf1 and, presumably, Upf2 and Upf3. Collectively, the results reported here and the earlier data supporting the *faux*-UTR model strongly suggest that inefficient termination is the trigger for NMD, possibly because the ribosome is transiently in a state that allows association of the Upf factors (He and Jacobson, 2015).

## STAR METHODS

### Yeast strains and plasmids

Yeast strains used in this study have the W303 background and are listed in the table of Key Resources. The wild-type (WT) strain and its isogenic *upf1Δ* derivative were described previously (He et al., 1997). Strains combining additional deletions with *upf1Δ*, i.e., *pbp1Δ upf1Δ*, *pab1Δ pbp1Δ upf1Δ*, and *pab1ΔC upf1Δ* were *upf1::KanMX6* derivatives of yDM146, yDM206, and YAS2239 (Kessler and Sachs, 1998; Mangus et al., 1998b) and were constructed by PCR-mediated strategies with pFA6a-KanMX6 as the template (Longtine et al., 1998). Fragments amplified by the primer pair upf1-KanMX6-F and upf1-KanMX6-R were transformed into respective yeast strains by the high-efficiency method (Schiestl and Gietz, 1989). Each of the genomic DNA deletions was confirmed by PCR analysis with the oligonucleotide pair upf1-SF and upf1-SR.

All of the plasmids and oligonucleotides used in this study are listed in Supplementary Tables 1 and 2. The plasmid pRS315-*LUC* was subcloned from plasmid YEplac181-*HA*-*LUC*-*SF* (Roy et al., 2015b) as follows: First, a PstI-XbaI fragment was isolated from YEplac181-*HA*-*LUC*-*SF* and cloned into pRS315 vector digested by PstI/XbaI. Then, a *TPI1 3’*-*UTR* fragment was amplified from YEplac181-*HA*-*LUC*-*SF* by the primer pair of TPI13U-F(XbaI)/TPI13U-R(NotI), digested by XbaI/NotI, and ligated into pRS315-*TPI1 promoter*-3X-*HA*-*LUC-StrepII-FLAG* digested by XbaI/NotI.

The pRS315-*LUC-PTCs* were generated by overlap PCRs using a common outside primer pair, TPI1-promoter-F (PstI)/TPI13U-R (NotI), and an internal primer pair that introduced PTCs at specific codons: codon 20 (LUC-PTC20-F/R), codon 41 (LUC-PTC41-F/R), codon 160 (LUC-PTC160-F/R), codon 386 (LUC-PTC386-F/R), codon 460 (LUC-PTC460-F/R), and codon 520 (LUC-PTC520-F/R). In each case, the final PCR products were digested by PstI/NotI and cloned into pRS315 digested by PstI/NotI.

pRS315 plasmids containing *LUC*-*PTC520*-Δ1-99, -Δ1-198, -Δ101-298, -Δ201-298, -Δ1-24, -Δ25-50, -Δ50, and -L alleles were constructed by direct PCR of specific *TPI1 3’* regions using primer pairs of TPI3U-100-F(XbaI)/TPI13U-R(NotI), TPI3U-200-F(XbaI)/TPI13U-R(NotI), TPI13U-F(XbaI)/ TPI3U-100-R(NotI), TPI13U-F(XbaI)/ TPI3U-200-R(NotI), TPI3U-25-F(XbaI)/TPI13U-R(NotI), TPI3U-2550-F(XbaI)/TPI13U-R(NotI), TPI3U-50-F(XbaI)/TPI13U-R(NotI), and 200L-F(XbaI)/TPI13U-R(NotI), respectively. In each case, the PCR fragment digested by XbaI/NotI was used to replace the *TPI1 3’* fragment in pRS315-*LUC*-*PTC520*. pRS315 plasmids containing *LUC*-*PTC520*-*S* and *LUC*-*PTC520*-*3U60* were generated by ligating the annealed DNA fragments of oligo pairs 100S-F/100S-R and GAL7-3U-F/GAL7-3U-R directly into pRS315-*LUC*-*PTC520* digested by XbaI/NotI, respectively.

pRS315 plasmids containing *LUC-PTC520-3U120, −3U200, −3U300, −3U400, −3U500*, and *-3U600* were all constructed by overlap PCRs in two steps. First, using a common outside primer pair (RS315-F-SalI/RS315-R-KasI) and a specific internal primer pair (3U120-F/3U120-R, 3U200-F/3U200-R, 3U300-F/3U300-R, 3U400-F/3U400-R, 3U500-F/3U500-R, and 3U600-F/3U600-R), we generated a set of pRS315 vectors with a *GAL7* polyadenylation signal (Bucheli et al., 2007) inserted at different locations of the 1 kb SalI-KasI fragment. Second, a PstI-XbaI fragment was isolated from pRS315-*LUC*-*PTC520* and ligated into this set of modified pRS315 vectors digested by PstI/XbaI. This set of *LUC*-*PTC520*-*xxUTR* alleles from pRS315 plasmids generates transcripts with 3’-UTR lengths ranging from 120 to 600 nt.

pRS316 plasmids containing *LUC* or *LUC*-*PTC* alleles were constructed in two steps. First, a HindIII-XbaI DNA fragment was isolated from the respective pRS315-*LUC* or *LUC*-*PTC* plasmids and cloned into pRS316. Then a XbaI-NotI fragment containing the *TPI1 3’* region was cloned downstream of the *LUC* or *LUC*-*PTC* ORF.

pRS315 derived plasmids were transformed into WT or *upf1Δ* yeast, and pRS316 derived plasmids were transformed into *pab1Δ pbp1Δ upf1Δ* or *pbp1Δ upf1Δ* yeast cells by the high-efficiency method (Schiestl and Gietz, 1989).

### Cell growth and western analysis

Cells were grown in 25 ml of synthetic complete (SC) media lacking leucine or uracil at 30°C to an optical density at 600nm (OD_600_) of 0.7. For each culture expressing a specific *LUC* or *LUC*-*PTC* allele, 10 OD_600_ units were harvested for western analysis and 4 OD_600_ units from the same culture were collected for RNA extraction. Cell pellets for western analyses were resuspended in 110 µl RIPA buffer (25 mM Tris-HCl pH 7.4, 150 mM NaCl, 1% NP-40, 1 mM EDTA, 5% glycerol) with 1mM PMSF and 1X protease inhibitor mixture (Thermo Scientific, #A32965). The cell suspensions were lysed by vortexing for 30 sec with 0.1g pre-washed glass beads (Sigma-Aldrich, #8772), followed by 30 sec on ice, for 8 cycles. The lysates were centrifuged at 12,000rpm for 15 min in a Sorvall Legend X1R centrifuge, at 4°C. Aliquots (90 µl) of the supernatants were mixed with Laemmli’s SDS-sample buffer (Boston BioProducts, #BP-110R) followed by boiling for 5 min. The samples were then resolved by 10% SDS-PAGE, transferred to Immobilon-P membrane (Millipore, #IPVH0010), and incubated with anti-HA antibodies (Sigma-Aldrich, #H3663, 1:2,000), anti-FLAG antibodies (Sigma-Aldrich, #F7425, 1:1,000 or SANTA CRUZ, #sc-51590, 1:100), or anti-Pgk1 antibodies (Thermo Fisher, #459250, 1:4,000) overnight at 4°C. The membrane was washed with PBST for 7 min, 3 times, then incubated with anti-Mouse secondary antibodies (Thermo Fisher, #A24506, 1:5,000-1:10,000) or anti-Rabbit secondary antibodies (Sigma-Aldrich, #GENA934, 1:5,000) for 45 min at RT. Then the membrane was washed with PBST for 7 min, 3 times. Proteins were detected using ECL reagents and Hyperfilm ECL (GE, #28906835). The signal was analyzed with MultiGauge software.

### RNA preparation and northern analysis

Total RNA from cell pellets was isolated as described previously (Herrick et al., 1990). Briefly, pellets were resuspended in 500 µl buffer A (50 mM NaOAc pH5.2, 10 mM EDTA, 1% SDS, 1% DEPC), extracted with 500 µl RNA-phenol (phenol saturated with 50 mM NaOAc pH5.2, 10 mM EDTA) by vortexing for 10 sec, followed by 50 sec incubation in a 65°C water bath, for 6 cycles. Tubes were centrifuged at 12,000rpm in an Eppendorf 5417C centrifuge for 10 min at RT. The aqueous layers were recovered and followed by another phenol extraction. Phenol/chloroform (500 µl) was added to the recovered aqueous layer, and the mixture was vortexed for 2 min at RT followed by 10 min centrifugation at 12,000rpm. The aqueous layers were recovered and subjected to another phenol/chloroform extraction. The RNAs were precipitated by adding 40 µl NaOAc (3 M, pH 5.2) and 1 ml ethanol and incubated for 2 hours at −80°C. Tubes were centrifuged at 12,000rpm for 15 min at 4°C. Pellets were washed with 70% ethanol and re-dissolved in 50 µl RNase free distilled water. Aliquots (15 µg) of each RNA sample were loaded onto a 1% agarose/formaldehyde/MOPS gel and electrophoresed, blotted, and hybridized as described previously (He and Jacobson, 1995). Random-primed DNA probes made from NcoI-XbaI *LUC* fragments were used to detect *HA-LUC-SF* mRNAs and full-length *SCR1* probes were used to detect the *SCR1* RNA (loading control). [α-^32^P]-dCTP (Perkin Elmer, Blu513Z) and a random primed DNA labeling kit (Roche, # 11-004-760-001) were used to generate probes according to the manufacturer’s protocol. Signals from northern blots were detected and analyzed by phosphorimaging using a Fujifilm bio-imaging analyzer (BAS-2500) and MultiGauge software.

### Calculation of readthrough and termination efficiencies

The following formulas were used to determine the relative efficiencies of readthrough and termination:

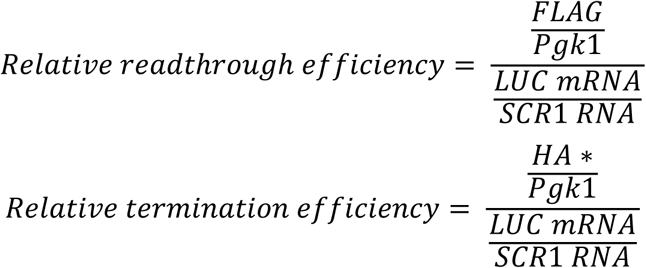

In these formulas *FLAG, Pgk1*, and *HA\** respectively represent the amounts of full-length FLAG-tagged Luc protein, Pgk1 protein, and prematurely terminated HA-tagged Luc protein determined by western blotting, and *LUC mRNA* and *SCR1 RNA* respectively designate the levels of these two transcripts determined by northern blotting. The results shown in the figures are the average of three independent experiments +/− SEM unless otherwise indicated.

## ACKNOWLEDGMENTS

This work was supported by grants to A.J. (5R01 GM27757-37 and 1R35GM122468-03) from the U.S. National Institutes of Health. We thank Robin Ganesan, Kotchaphorn Mangkalaphiban, Andrei Korostelev, Christine Carbone, and Denis Susorov for helpful discussions and insightful comments about the manuscript.

## AUTHOR CONTRIBUTIONS

C.W., B.R., F.H., and A.J. conceived and designed the experiments, C.W. and B.R. carried out the experiments, C.W. and A.J. wrote the paper with input from all authors, and A.J. obtained funding for the study.

## KEY RESOURCES TABLE

| REAGENT or RESOURCE | SOURCE | IDENTIFIER |
| --- | --- | --- |
| Antibodies |  |  |
| Rabbit polyclonal anti-FLAG | Sigma-Aldrich | Cat#F7425 |
| Mouse monoclonal anti-OctA-Probe (F-tag-01) | SANTA CRUZ | Cat#sc-51590 |
| Mouse monoclonal anti-Pgk1 (22C5D8) | Thermo Fisher | Cat#459250 |
| Donkey polyclonal anti-Mouse Secondary | Thermo Fisher | Cat#A24506 |
| Donkey anti-Rabbit Secondary | Sigma-Aldrich | Cat#GENA934 |
| Mouse monoclonal anti-HA | Sigma-Aldrich | Cat#H3663 |
| Chemicals, Peptides, and Recombinant Proteins |  |  |
| Protease inhibitor tablets, EDTA-free | Thermo Fisher | Cat#A32965 |
| dCTP, [α-<sup>32</sup>P]-6000Ci/mmol | Perkin Elmer | Cat#BLU513Z |
| MG-132 | Sigma-Aldrich | Cat#474790 |
| Synthetic HA-Luc19:<br>**MGYPYDVPDYAGYPYDVPDYAGSYPYDVPDYA**<br>**MEDAKNIKKGPAPFYPLED** | GenScript | This paper |
| Critical Commercial Assays |  |  |
| Random Primed DNA Labeling Kit | Sigma-Aldrich | Cat#11004760001 |
| Experimental Models: Organisms/Strains |  |  |
| *S. cerevisiae*: *upf1*Δ (HFY871): *MATa ade2-1 his3-11,15 leu2-3,112 trp1-1 ura3-1 can1-100 upf1::HIS3* | Jacobson lab | He et al., 1997 |
| *S. cerevisiae*: WT (HFY114): *MATa ade2-1 his3-11,15 leu2-3,112 trp1-1 ura3-1 can1-100* | Jacobson lab | He et al., 1997 |
| *S. cerevisiae*: *pab1*Δ *pbp1*Δ *upf1*Δ: *MATa ade2-1 his3-11,15 leu2-3,112 trp1-1 ura3-1 can1-100 pab1::HIS3 pbp1::LEU2 upf1::KanMX6* | Jacobson lab | This paper |
| *S. cerevisiae*: *pab1*ΔC *upf1*Δ: *MATa ade2-1 his3-11,15 leu2-3,112 trp1-1 ura3-1 can1-100 pab1*ΔC::HIS3 *upf1::KanMX6* | Jacobson lab | This paper |
| *S. cerevisiae*: *pbp1*Δ *upf1*Δ: *MATa ade2-1 his3-11,15 leu2-3,112 trp1-1 ura3-1 can1-100 pbp1::LEU2 upf1::KanMX6* | Jacobson lab | This paper |
| Software and Algorithms |  |  |
| MultiGauge software | Fujifilm | Science lab 2005 |

---

**Figure S1.**
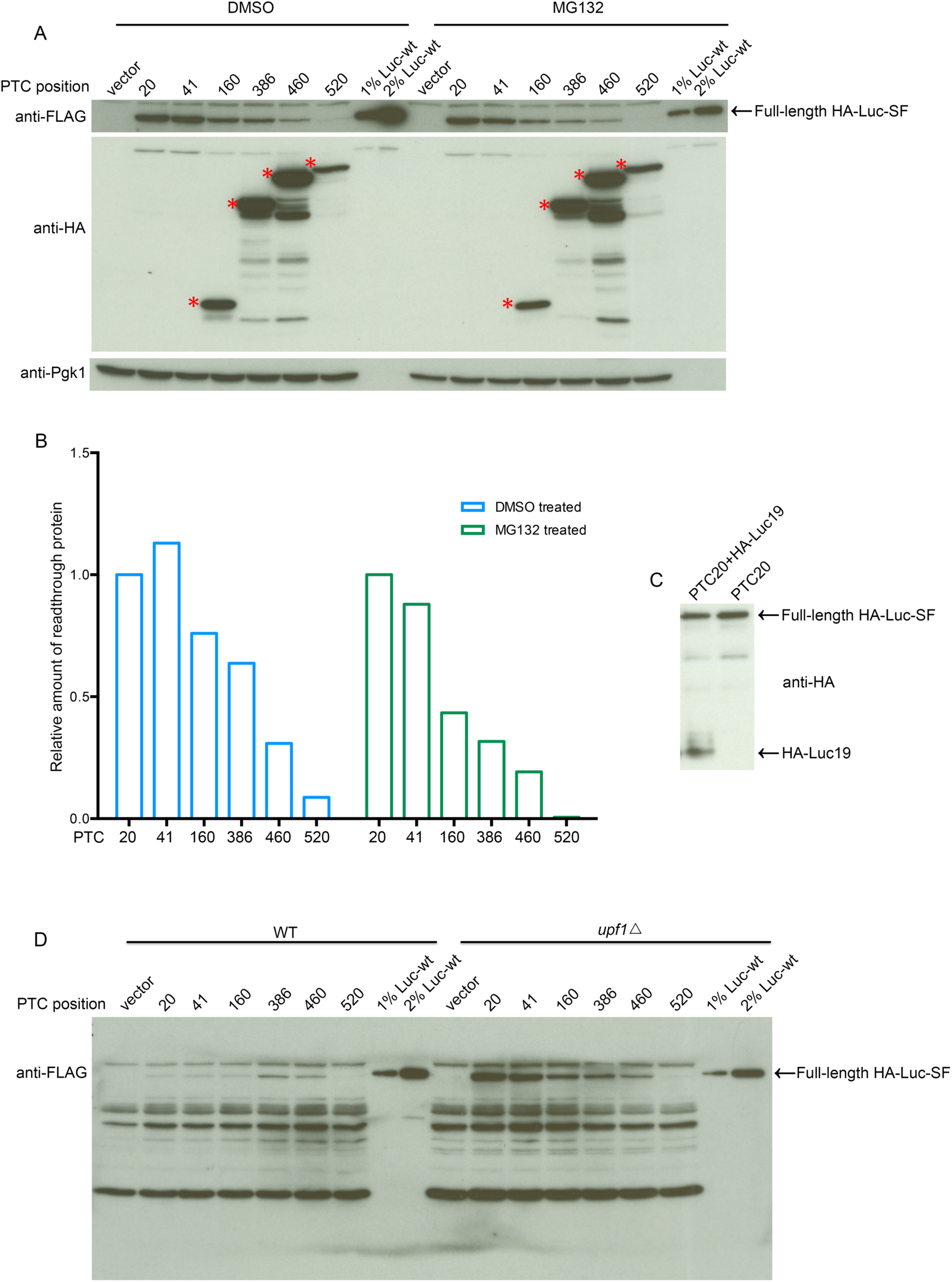
MG132 treatment has no significant effect on the expression of readthrough and premature termination products generated from *LUC* PTC alleles in *upf1Δ* cells. **A. Western analyses of polypeptides expressed from different *LUC* PTC alleles in mock- or MG132-treated *upf1Δ* cells**. Each of the cultures was processed as in Figure 1B, except that the growth media were supplemented with either 0.05% DMSO or 0.1mM MG132. **B. Relative levels of full-length readthrough products expressed from each of the six LUC PTC alleles in mock- or MG132-treated *upf1Δ* cells.** Densitometry was utilized to quantitate the western blots in part A and relative recoveries were calculated by determining each sample’s full-length Luc protein level (anti-FLAG band) normalized to its Pgk1 level. The *LUC* PTC20 sample was set as 1 in each group. **C. Western analysis of the HA-Luc19 peptide.** Synthetic HA-Luc19 peptide was mixed with one of two aliquots of *LUC* PTC20 cell lysate and both samples were processed for western blotting, with detection by anti-HA antibodies. Left lane: *LUC* PTC20 cell lysate containing HA-Luc19 peptide; right lane: *LUC* PTC20 cell lysate only. **D. Complete western blot used to depict full-length readthrough protein in the top panel of Figure 1B.** The same western blot from which a segment was used to illustrate the extent of readthrough protein accumulation in Figure 1B is shown in its entirety.

**Figure S2.**
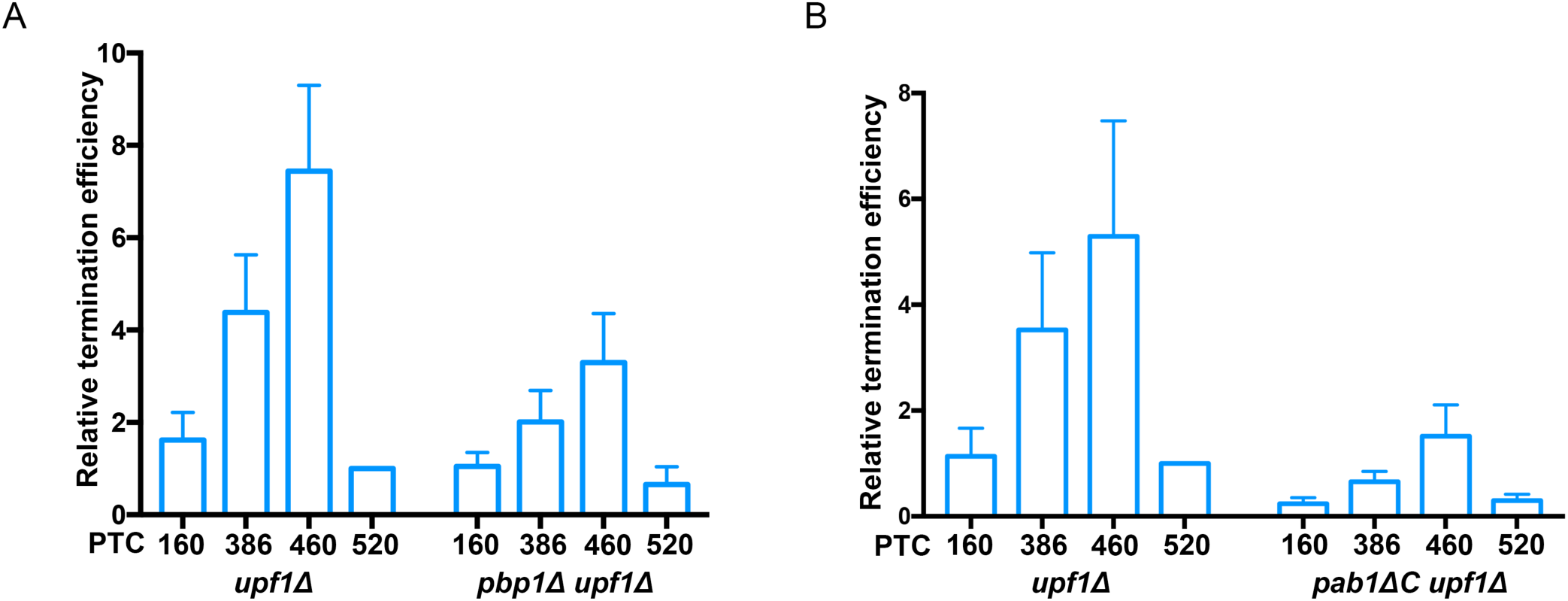
Relative termination efficiencies of the different *LUC* PTC alleles. **A. Relative termination efficiencies of *LUC* PTCs in *upf1Δ* or *pbp1Δ upf1Δ* cells.** Densitometry and phosphorimaging were respectively utilized to quantitate the western blots of Figure 6A and the northern blots of Figure 6B. Relative termination efficiencies were calculated as the level of prematurely terminated Luc protein level (anti-HA) normalized to the level of Pgk1 divided by the *LUC* mRNA level normalized to the *SCR1* level. The efficiency of termination for the *LUC* PTC520 allele in *upf1Δ* cells was set as 1. **B. Relative termination efficiencies of *LUC* PTCs in *upf1Δ* or *pab1ΔC upf1Δ* cells.** The western and northern blots of Figure 7A and B were used to calculate relative termination efficiencies as in part A. Three independent experiments were performed and the results shown are the average of all three +/− SEM.

**Supplementary Table 1.** Plasmids used in this study

| Name | Description |
| --- | --- |
| pRS315 | Yeast centromere-based vector with *LEU2* selection marker |
| pRS315-*LUC* | Contains *TPI1* promoter, N-terminal 3XHA tag, firefly luciferase gene, C-terminal StreptII-FLAG (SF) tag, and a 298-bp *TPI1* 3'-UTR fragment |
| pRS315-*LUC-PTC20* | Same as pRS315-*LUC* but contains TGA CAA at *LUC* codon position 20 and 21. |
| pRS315-*LUC-PTC41* | Same as pRS315-*LUC* but contains TGA CAA at *LUC* codon position 41 and 42. |
| pRS315-*LUC-PTC160* | Same as pRS315-*LUC* but contains TGA CAA at *LUC* codon position 160 and 161. |
| pRS315-*LUC-PTC386* | Same as pRS315-*LUC* but contains TGA CAA at *LUC* codon position 386 and 387. |
| pRS315-*LUC-PTC460* | Same as pRS315-*LUC* but contains TGA CAA at *LUC* codon position 460 and 461. |
| pRS315-*LUC-PTC520* | Same as pRS315-*LUC* but contains TGA CAA at *LUC* codon position 520 and 521. |
| pRS316 | Yeast centromere-based vector with *URA3* selection marker |
| pRS316-*LUC* | Contains *TPI1* promoter, N-terminal 3XHA tag, firefly luciferase gene, C-terminal StreptII-FLAG (SF) tag, and a 298-bp *TPI1* 3'-UTR fragment |
| pRS316-*LUC-PTC20* | Same as pRS316-*LUC* but contains TGA CAA at *LUC* codon position 20 and 21, C-terminal StreptII-FLAG (SF) tag, *TPI1* 3'-UTR. |
| pRS316-*LUC-PTC41* | Same as pRS316-*LUC* but contains TGA CAA at *LUC* codon position 41 and 42. |
| pRS316-*LUC-PTC160* | Same as pRS316-*LUC* but contains TGA CAA at *LUC* codon position 160 and 161. |
| pRS316-*LUC-PTC386* | Same as pRS316-*LUC* but contains TGA CAA at *LUC* codon position 386 and 387. |
| pRS316-*LUC-PTC460* | Same as pRS316-*LUC* but contains TGA CAA at *LUC* codon position 460 and 461. |
| pRS316-*LUC-PTC520* | Same as pRS316-*LUC* but contains TGA CAA at *LUC* codon position 520 and 521. |
| pRS315-*LUC-PTC520-Δ1-99* | Same as pRS315-*LUC-PTC520* but contains a 99-bp deletion from position 1 to 99 in the *TPI1* 3'-UTR fragment. |
| pRS315-*LUC-PTC520-Δ1-198* | Same as pRS315-*LUC-PTC520* but contains a 198-bp deletion from position 1 to 198 in the *TPI1* 3'-UTR fragment |
| pRS315-*LUC-PTC520-Δ101-298* | Same as pRS315-*LUC-PTC520* but contains a 198-bp deletion from position 101 to 298 in the *TPI1* 3'-UTR fragment. |
| pRS315-*LUC-PTC520-Δ201-298* | Same as pRS315-*LUC-PTC520* but contains a 98-bp deletion from position 201 to 298 in the *TPI1* 3'-UTR fragment. |
| pRS315-*LUC-PTC520-Δ1-24* | Same as pRS315-*LUC-PTC520* but contains a 24-bp deletion from position 1 to 24 in the *TPI1* 3'-UTR fragment. |
| pRS315-*LUC-PTC520-Δ25-50* | Same as pRS315-*LUC-PTC520* but contains a 25-bp deletion from position 25 to 50 in the *TPI1* 3'-UTR fragment. |
| pRS315-*LUC-PTC520-Δ1-50* | Same as pRS315-*LUC-PTC520* but contains a 50-bp deletion from position 1 to 50 in the *TPI1* 3'-UTR fragment. |
| pRS315-*LUC-PTC520-ΔFL* | Same as pRS315-*LUC-PTC520* but lacks the 298-bp *TPI1* 3'-UTR fragment. |
| pRS315-*LUC-PTC520*-3U60 | Same as pRS315-*LUC-PTC520-ΔFL* but contains a 60-bp insertion of the *GAL7* poly(A) adenylation signal immediately downstream the *LUC* stop codon |
| pRS315-*LUC-PTC520*-3U120 | Same as pRS315-*LUC-PTC520-ΔFL* but contains a 60-bp insertion of the *GAL7* poly(A) adenylation signal 60 nts downstream the *LUC* stop codon in the vector sequence. |
| pRS315-*LUC-PTC520*-3U200 | Same as pRS315-*LUC-PTC520-ΔFL* but contains a 60-bp insertion of the *GAL7* poly(A) adenylation signal 140 nts downstream the *LUC* stop codon in the vector sequence. |
| pRS315-*LUC-PTC520*-3U300 | Same as pRS315-*LUC-PTC520-ΔFL* but contains a 60-bp insertion of the *GAL7* poly(A) adenylation signal d 240 nts downstream the *LUC* stop codon in the vector sequence. |
| pRS315-*LUC-PTC520*-3U400 | Same as pRS315-*LUC-PTC520-ΔFL* but contains a 60-bp insertion of the *GAL7* poly(A) adenylation signal 340 nts downstream the *LUC* stop codon in the vector sequence. |
| pRS315-*LUC-PTC520*-3U500 | Same as pRS315-*LUC-PTC520-ΔFL* but contains a 60-bp insertion of the *GAL7* poly(A) adenylation signal 440 nts downstream the *LUC* stop codon in the vector sequence. |
| pRS315-*LUC-PTC520*-3U600 | Same as pRS315-*LUC-PTC520-ΔFL* but contains a 60-bp insertion of the *GAL7* poly(A) adenylation signal 540 nts downstream the *LUC* stop codon in the vector sequence. |
| pRS315-*LUC-PTC520-S* | Contains *TPI1* promoter, N-terminal 3XHA tag, firefly luciferase gene with TGA CAA at *LUC* codon position 520 and 521, C-terminal StrepII-FLAG (SF) tag, and a *TPI1* 3'-UTR fragment from position 100 to 199 with the *GAL7* poly(A) adenylation signal inserted at position 130. |
| pRS315-*LUC-PTC520-L* | Contains *TPI1* promoter, N-terminal 3XHA tag, firefly luciferase gene with TGA CAA at *LUC* codon position 520 and 521, C-terminal StrepII-FLAG (SF) tag, and a *TPI1* 3'-UTR from position 100 to 298 with the *GAL7* poly(A) signal inserted position 130. |

**Supplementary Table 2.**
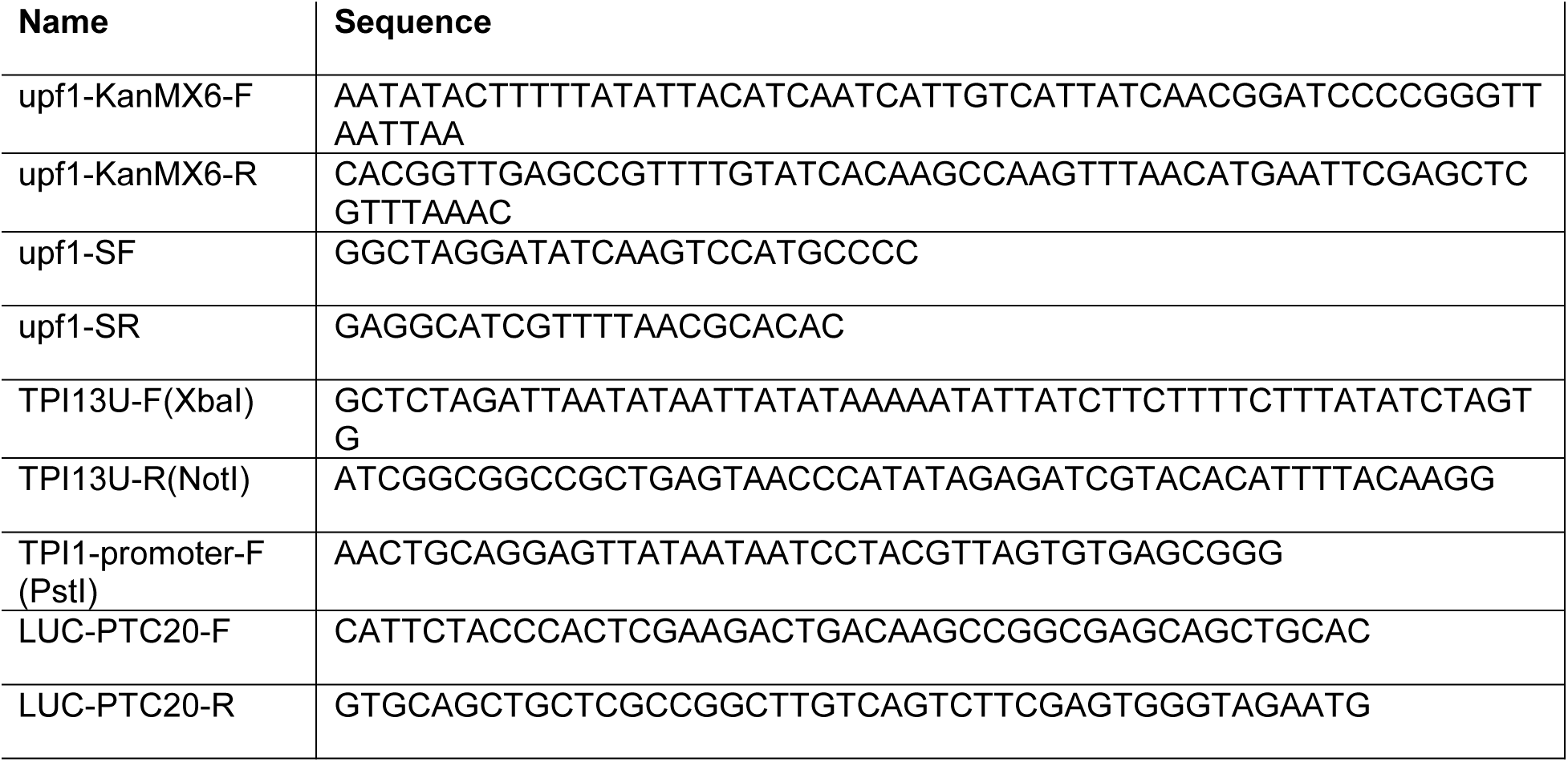
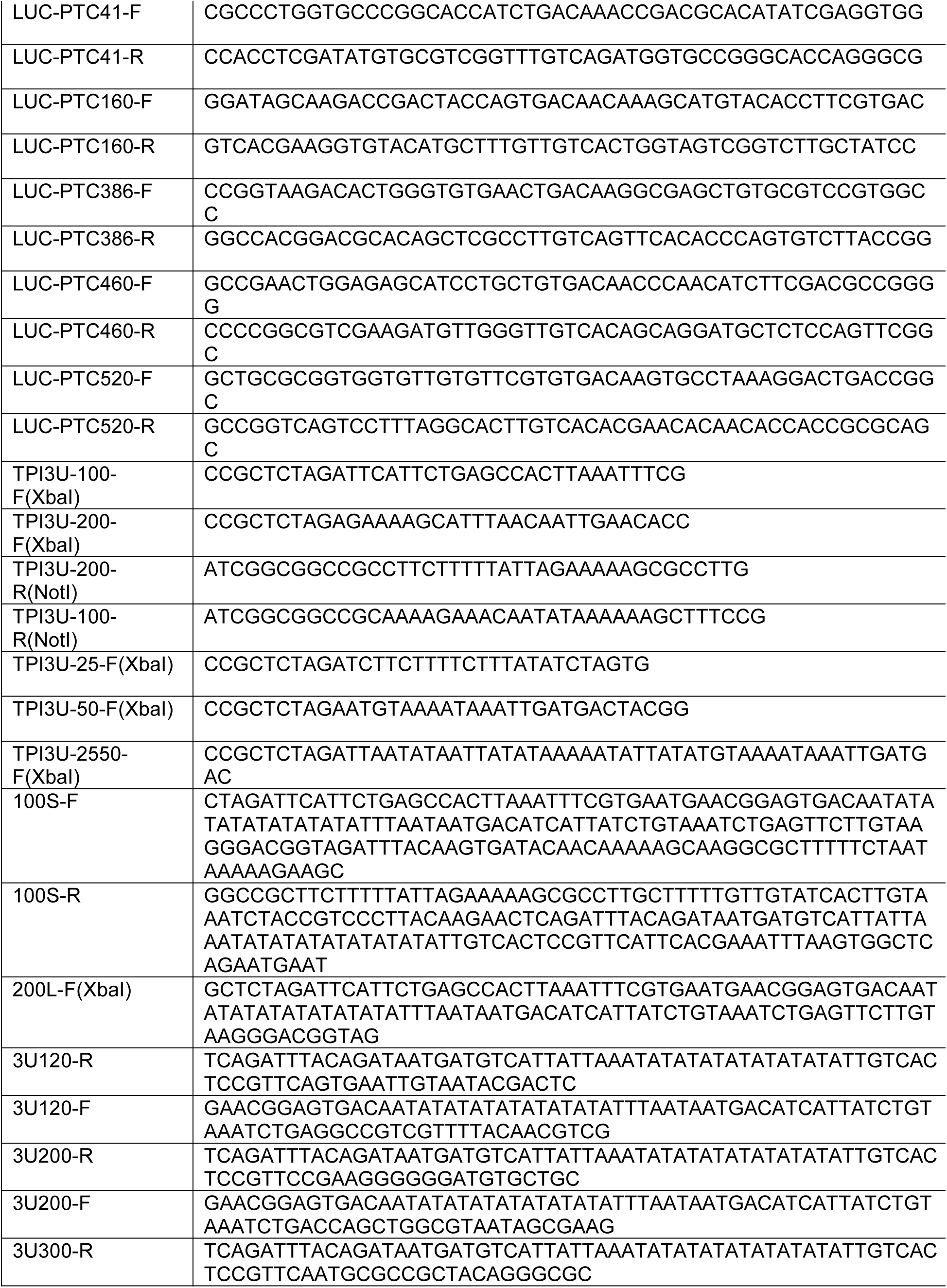
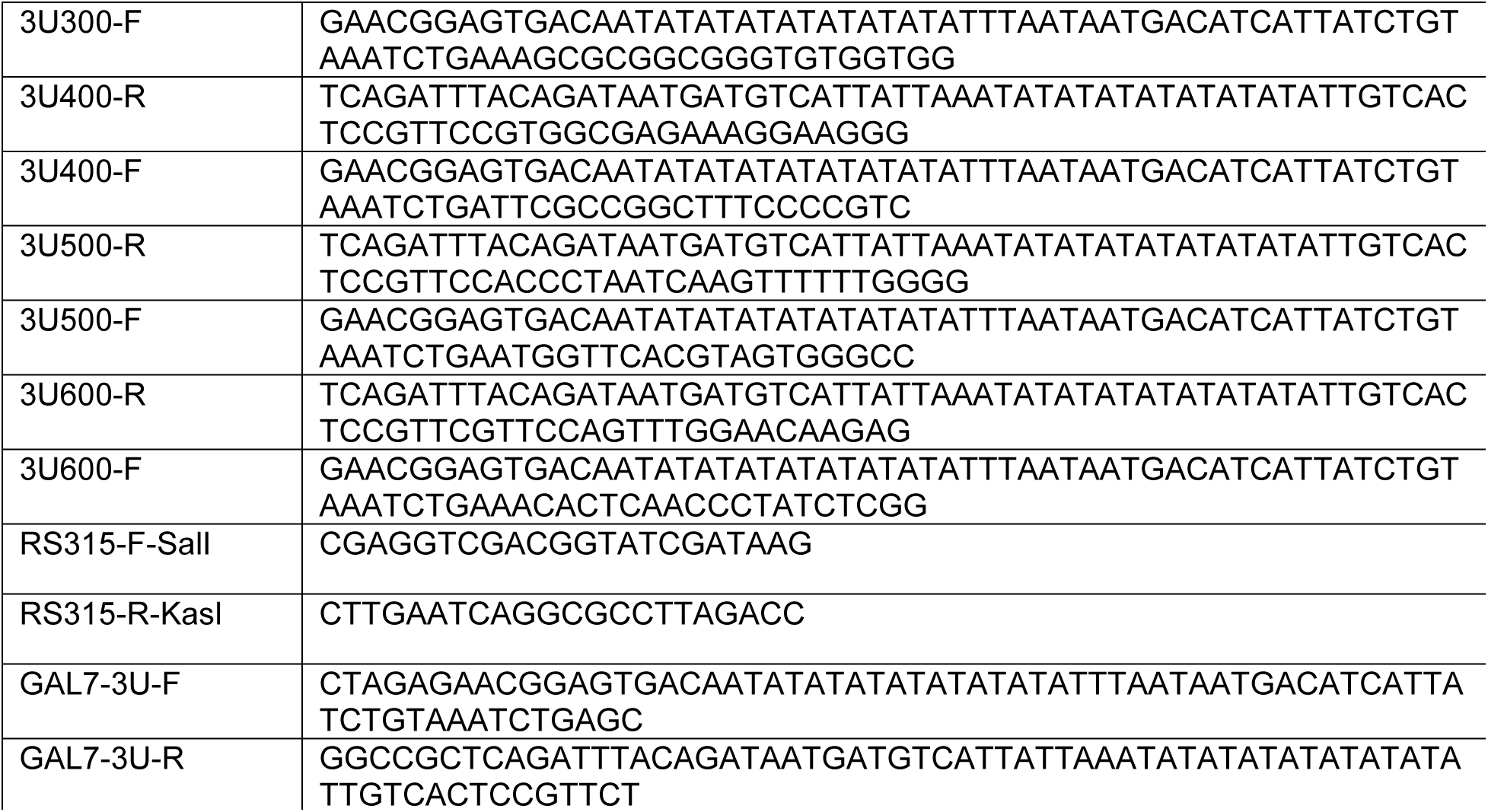
Oligonucleotides used in this study

